# Structural capture of an intermediate transport state of a CLC CI^-^/H^+^ antiporter

**DOI:** 10.1101/384404

**Authors:** Kunwoong Park, Hyun-Ho Lim

## Abstract

The CLC family proteins are involved in a variety of cellular processes, where chloride homeostasis needs to be controlled. Two distinct classes of CLC proteins, Cl^-^ channels and Cl^-^/H^+^ antiporters, have been functionally and structurally investigated over the last several decades. Recent studies have revealed that the conformational heterogeneity of the critical glutamate residue, Glu_ex_ could explain the transport cycle of CLC-type Cl^-^/H+ antiporters. However, the presence of multiple conformations of the Glu_ex_ has been suggested from combined structural snapshots of two different CLC antiporters. Thus, we aimed to investigate the presence of these three intermediate conformations in CLC-ec1, the most deeply studied CLC at both functional and structural levels. By comparing crystal structures of E148D, E148A mutant and wildtype CLC-ec1 with varying anion concentrations, we suggest that the Glu_ex_ indeed take at least three distinct conformational states in a single CLC antiporter, CLC-ec1.

## INTRODUCTION

The CLC family proteins are evolutionarily well conserved proteins expressed in the biological membranes and are critical for diverse physiological processes that control Cl^-^concentration including the regulation of skeletal muscle and neuronal excitability, acidification of intracellular organelles, and cell volume regulation, among others^1-3^. Genetic mutations of CLC genes are linked to a variety of pathological outcomes, such as myotonia, leukodystrophy, deafness, retinal degeneration, Bartter’s syndrome, Dent’s disease, and osteopetrosis^4-6^. The CLC family proteins are classified functionally into two mechanistically distinct protein families of Cl^-^channels and Cl^-^/H^+^ antiporters. The CLC-type Cl^-^ channels allow Cl^-^ to move across the membrane according to its electrochemical gradient. In contrast, CLC-type Cl^-^/H^+^ antiporters catalyze energetically *uphill* movement of Cl^-^ with the compensation of coupled *downhill* movement of H^+^, or vice versa^7-9^. In addition to conventional Cl^-^/H^+^ antiporters, the plant system is equipped with CLC-type NO_3_^-^/H^+^ antiporters that transport nitrate into the vacuoles^10,11^ and some bacteria use CLC-type F^-^/H^+^ antiporters to resist the environmental toxicity of fluoride^12,13^.

Regardless of their mechanistic diversity, the CLC family proteins share a highly conserved key glutamate residue, called the external glutamate (Glu_ex_) or the gating glutamate (Glu_gate_), which serves as a gate for the Cl^-^-conducting pathway in both CLC channels and antiporters as well as for H^+^ transit of CLC antiporters during the transport cycle^8,9,14^. Structural studies of various CLC proteins from bacteria to mammals have revealed their molecular architectures at atomic resolution^15-22^, which immediately suggest the molecular mechanisms of how CLC proteins transport ions. In the structures, the Glu_ex_ adopts at least three different conformations along the Cl^-^ transport pathway: *Up-, Middle-*, or Down-conformation (Figure 1). Based on the rotamerically distinct positions of the Glu_ex_ in the CLC structures, it was suggested that the glutamate side chain competes for two anion binding sites with permeant chloride ions and accepts a proton after expelling two chloride ions during the transport cycle of Cl^-^ and H^+^ exchange^19^.

**Figure 1.**
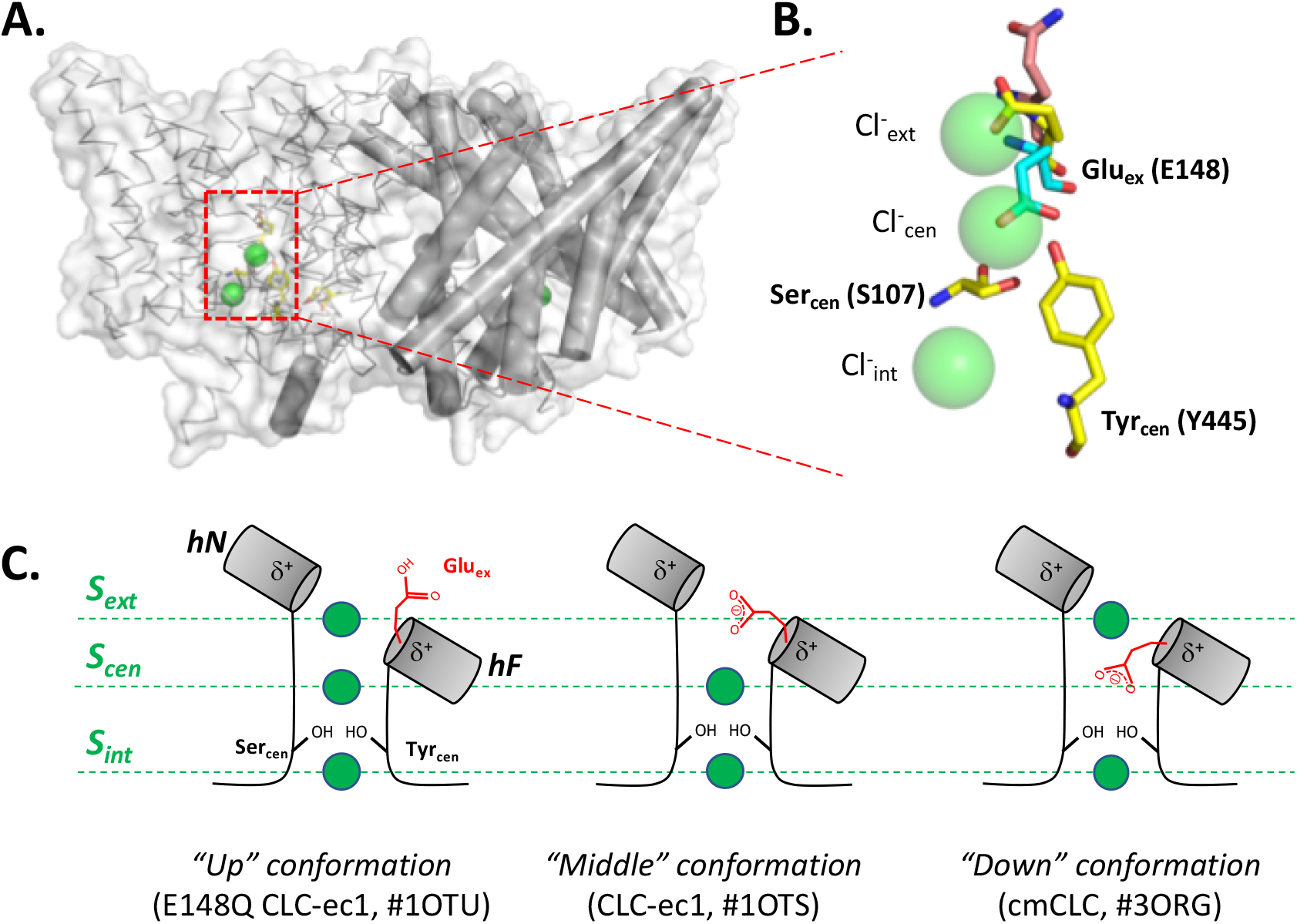
Multiple conformations of the Glu_ex_ residue in the transport cycle of CLC antiporters with known structures. A. Overall structure of CLC-ec1. Two protomers are presented as either *ribbon* or *cylinder* models. Key residues (Glul48, Glu203, SerlO7, and Tyr445) involved in Cl^-^/H^+^ transport are represented as stick models. The *red* dashed box indicates the region depicted in B. B. Conformations of CLC antiporters near the Cl^-^ binding site. Critical residues(Glu_ex_, Ser_cen_, and Tyr_cen_) for Cl^-^ transport in the structures of CLC antiporters are shown as stick models (*yellow*, carbon in wild-type CLC-ec1; *magenta*, carbon in E148Q CLC-ec1; *cyan*, carbon in cmCLC for carbon; *blue*, nitrogen; and *red*, oxygen). Chloride ions are shown as *green* spheres. The PDB accession numbers for wild-type CLC-ec1, E148Q CLC-ec1, and cmCLC are #IOTS, #10TU, and #3ORG, respectively. C. Diagram of conformational diversity of Glu_ex_ in CLC antiporters. The internal, central and external anion binding sites are marked as *S_ext_, S_cen_*, and *S*_int_, respectively. Note that helix F (*hF*) and helix N (*hN*) offer helical dipoles (δ^+^) to stabilize both Cl^-^ and Glu_ex_ binding at S_ext_^15^

Though three alternative Glu^ex^ conformations would explain 2:1 coupled movement of Cl^-^ and H^+^ in Cl^-^/H^+^ antiporters, no single CLC protein has shown three conformational states structurally: *Up-* and *Middle*-conformations have been observed in the deeply studied CLC-ec1 from *E.coli*^15,16^; *Down*-conformation in a eukaryotic CLC, cmCLC from a thermophilic red alga^17^; and *Up-* and *Down*-conformations in a distinct CLC-type F^-^/H^+^ antiporter clade, CLC-Eca^22^. Thus, whether the Glu_ex_ in a single CLC protein adopts all three rotameric conformations during the transport cycle remains unclear.

Recent studies on the mutation of the external glutamate (Glu_ex_) in CLC-type Cl^-^/H^+^ antiporters to aspartate, an amino acid with a methylene (-CH2 group) shorter and a similar pKa value, reported extremely slow ion transport in both CLC-ec1 and cmCLC^19^. Moreover, a mutation at the corresponding glutamate (E166D) in the CLC-0 channel reduced single-channel conductance as well as drastically decreased open probability^23,24^. These previous observations support the idea that the Glu_ex_ has three conformational states *(Up-, Middle-*, and Down-conformations; Fig. 1C), and reflect the penalty of the shorter aspartate side chain to reach the *S_cen_*. Recently, it was also suggested that CLC-ec1 could adopt a cmCLC-like state by a conformational locking of the Glu_ex_ at the central anion binding site (*S_cen_*), which is induced by high external Cl^-^ concentration^25^. Computational study also proposed a transport cycle in which the Glu_ex_ could occupy the **S_cen_** in CLC-ec1^26^.

However, the presence of cmCLC-like “*Down*“ conformation, the Glu_ex_ occupying the **S_cen_**, has not been observed in CLC-ec1 structure, only in cmCLC. Thus, we aimed to tackle the following: (1) elucidating the structural change in the E148D mutant CLC-ec1, and explain the mechanistical consequences of E148D mutation, if it exists, (2) visualizing the structural transition occurring due to Asp148 protonation of the E148D mutant by comparing its structure with E148N mutant, and (3) obtaining structural insight into the conformational changes of the Glu_ex_, and especially into its presence at the **S_cen_** in CLC-ec1 during the transport cycle.

Structural and functional examinations of E148D and E148N CLC-ec1 revealed unexpected rotameric change upon protonation/deprotonation of Asp148, which resulted in limited solution accessibility to the Asp148 residue and could slow ion transport. From the comparison of the E148D mutant and halide-free wild-type structures, we provide evidence for the presence of a new intermediate “*Mid-low*” state in the transition between “*Middle*” and “*Down*” conformations. This “*Mid-low*” conformation of Glu_ex_ is enough to expel a chloride ion from the central binding site similarly observed in the structure of cmCLC. Additionally, we reconsolidated a previous observation^19^ with anomalously detectable short carboxylic acid, bromoacetate, which favors to bind in a *carboxylate-down* configuration to the ungated E148A mutant in the analogy of “*Down*” conformation of Glu_ex_. These results suggest that the external glutamate in a single CLC-type Cl^-^/H^+^ antiporter can undergo a structural twist from *Up-* to *Down*-conformation to exchange both Cl^-^ and H^+^.

## RESULTS

### Ion transport and anion binding in the E148D and E148N mutants

The E148D mutant mediates both Cl^-^ and H^+^ movements across reconstituted proteoliposomes with ~10-fold smaller transport rates (~180 Cl^-^/sec at pH 4.5) than that of wild-type CLC-ec1^19,27,28^ (Fig. 2A, B; Supplementary Table 1). The E148N mutant transports Cl^-^ with a turnover rate of ~110 Cl^-^/sec at pH 4.5, but fails to transport H^+^, as similarly observed in the E148Q mutant, a protonated Glu_ex_ mimic of wild-type CLC-ec1^29^ (Fig. 2A,B; Supplementary Table 1). Previous studies from several labs have indicated that Glu_ex_ is central to the pH- and voltage-dependent activation of both CLC-type channels and antiporters^7,16,24,30-32^, so we accessed the effect of E148D mutation on pH-dependent activation of CLC-ec1. The E148D mutant showed two times faster transport at pH 4 than at pH 6, though pH-dependency was shallower than that of wild-type CLC-ec1(Supplementary Fig.1; Supplementary Table 1) ^30,32^.

**Figure 2:**
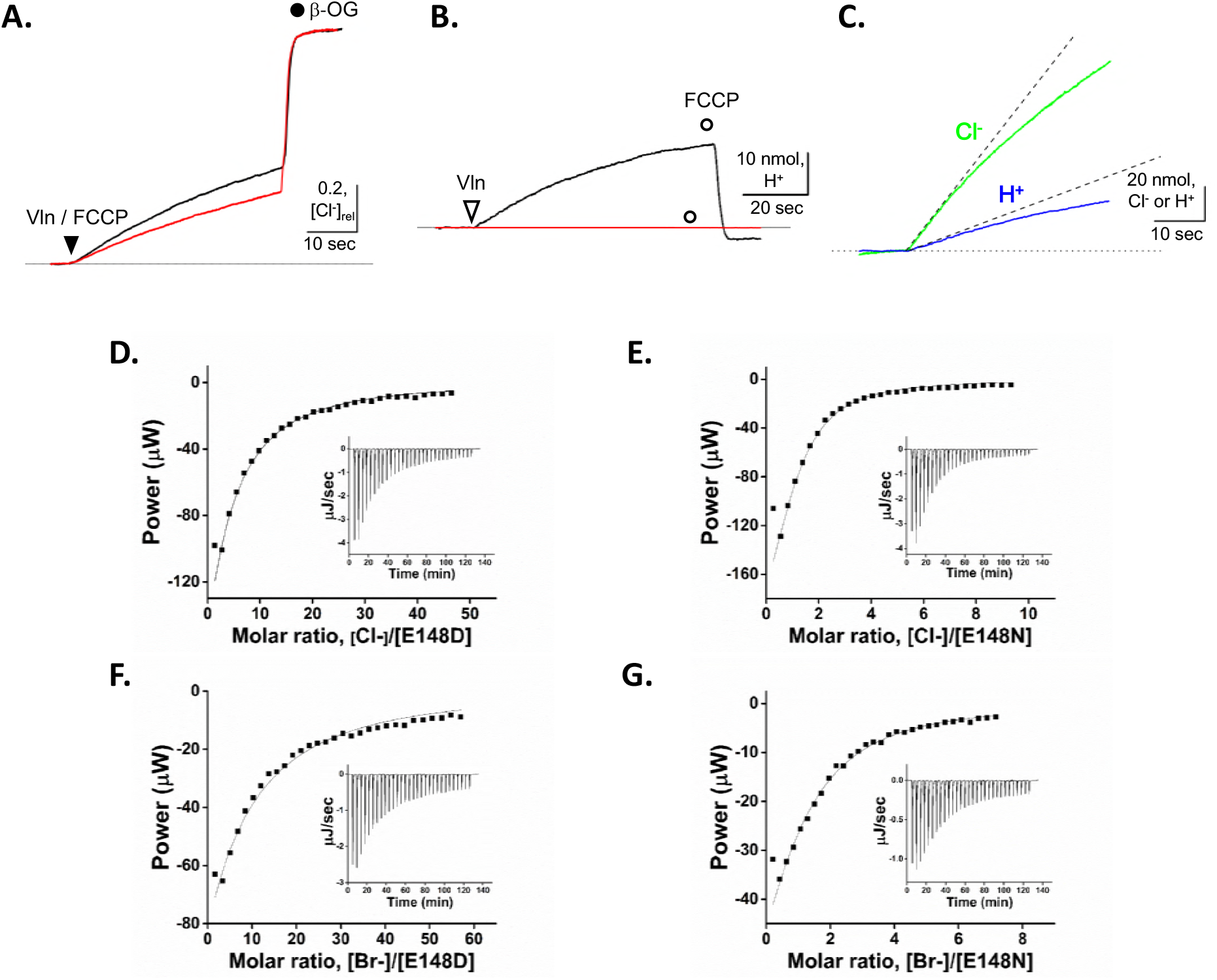
Ion transport and equilibrium anion binding affinities of E148D and E148N mutants. A.Representative data from Cl^-^ dump of proteoliposomes reconstituted with E148D *(black solid line)* and E148N *(reel solid line)* CLC-ec1. Cl^-^ transport is triggered by valinomycin (Vln) and FCCP *(arrow head)* and terminated by disrupting proteoliposomes with the detergent β-octylgluconate (β-OG*.filled circle)*. The dotted lines indicate baseline transport. B. Chloride-driven H^+^ transport of E148D *(black solid line)* andEl48N *(red solid line)* mutant. C. Coupled movement of Cl^-^ and H^+^ through the E148D mutant. *Green* and *blue* lines indicate concentration changes of Cl^-^ and H^+^, respectively, and dashed lines represent slopes for initial transports. Isothermal titration calorimetric measurements for equilibrium binding of Cl^-^ or Br^-^to E148D mutant (D and F, respectively) and E148N mutant CLC-ec1 (E and G, respectively). Solid curves indicate K_D_ values for Cl^-^ or Br^-^ binding to each protein. (*Insets*) Representative raw data for each experiment.

Interestingly, the stoichiometry of Cl^-^ and H^+^ transport in the E148D mutant is 3.8 ±0.2 (Fig. 2C, n=3) compared to ~2 to 1 in the wild-type^7,28^. Altered Cl^-^/H^+^ coupling ratios have been previously observed in mutants showing reduced Cl^-^ binding at the central binding site including Y445F and Y445I mutants^7,27^, and mutants limiting intracellular proton access such as E203K and E202L mutants^28,32^. Since altered Cl^-^/H^+^ coupling of these mutants showed increased Cl^-^ slippage owing to reduced H^+^ transport, E148D mutation could also affect H^+^ transport and change the stoichiometry of coupling.

In order to test whether markedly reduced ion transport observed in the E148D mutant is caused by changed anion binding affinity, we measured the equilibrium anion binding affinity of the E148D mutant by using isothermal titration calorimetry(ITC) at pH 7.5^28,29^.The E148D mutant has a slightly reduced Cl^-^ and Br^-^ binding affinities with the K_D_ of ~3.1 mM and ~5.1 mM, respectively, which are 2~4-fold weaker than those previously reported in wild-type CLC-ec1^28,29,33^ (Fig. 2D, F and Supplementary Table 2). The E148N mutant binds Cl^-^ with a K_D_ of ~0.1 mM, similarly to the E148Q mutant (~0.07mM)^29^, and binds Br^-^ with Kd of ~0.3 mM (Fig. 2E, G and Supplementary Table 2).

Although the Cl^-^ binding affinity is lowered in the E148D mutant (~4-fold), the binding affinity change cannot solely account for the slowed ion transport (~10-fold). On the other hand, the similar Cl^-^ binding affinity and marginal difference between Cl^-^ transport rates in E148N and E148Q mutants (~130 Cl^-^/sec and ~300 Cl^-^/sec, respectively; Supplementary Table 1, 2) were observed. These results strongly suggest that the conformational changes of Asp148 at the Glu_ex_ position and anion binding configuration in E148D mutants should differ from those of wild-type during the transport cycle. If this is the case, we should be able to see the structural changes induced by E148D mutation.

### Crystal structures of the external gate mutants E148D and E148N

To examine whether E148D mutation induces any structural change, we solved crystal structures of E148D and E148N mutants with F_AB_ antibody fragments^16^ at resolutions of 2.95Å and 2.7Å, respectively (Table 1). The overall structures of both E148D and E148N mutants are essentially identical to that of the wild-type (Ca r.m.s. deviation 0.26 and 0.19 Å, respectively; Supplementary Fig 2). Crystal structures were obtained with Br^-^ to take advantage of anomalous diffraction in identifying anion binding sites with confidence (Fig. 3, Supplementary Table 3). Since the binding affinity differences between Cl^-^ and Br^-^ in both E148D and E148N mutants are minimal (1.5~3-fold), Cl^-^ binding in these two mutants can be inferred from anomalous Br^-^ signals.

**Figure 3.**
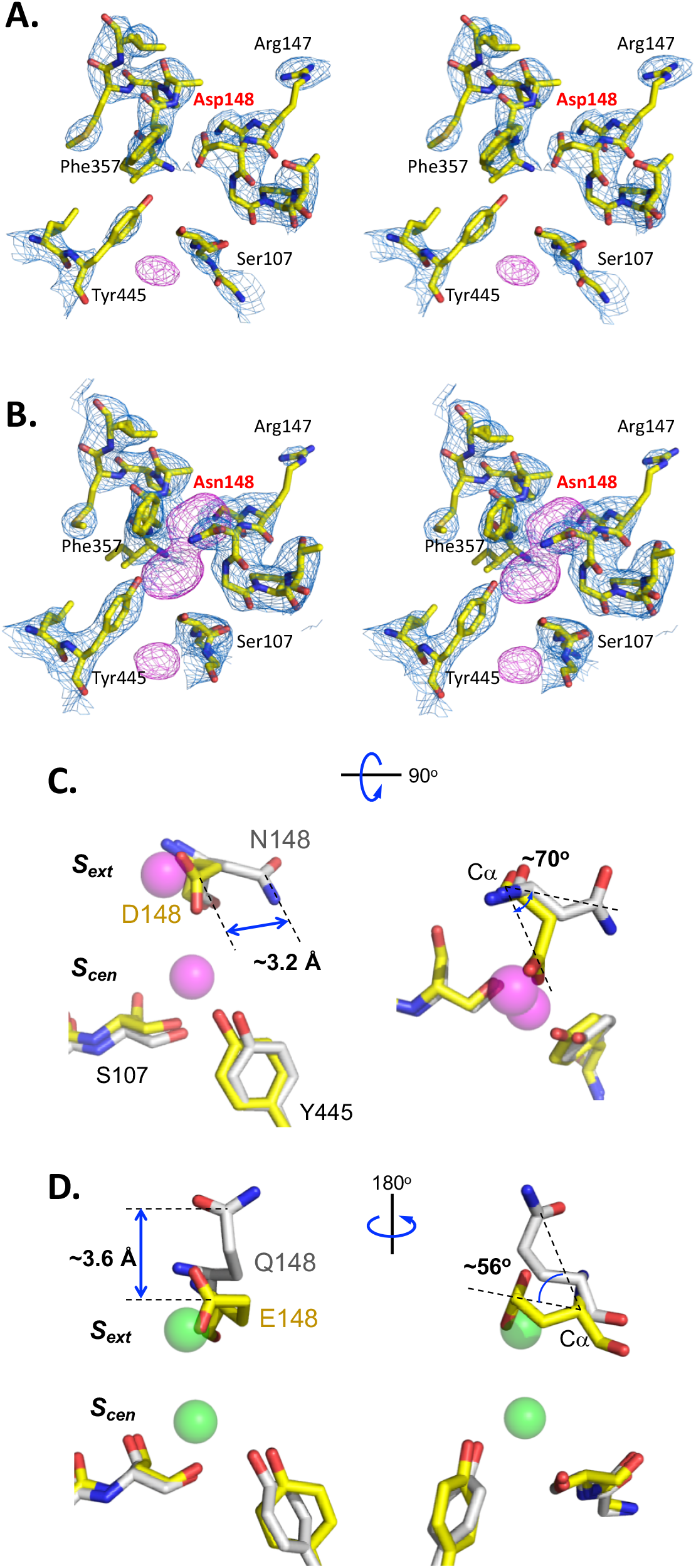
Crystal structures of E148D and E148N mutants near the Cl binding sites. Stereo-view structures of E148D (A) and E148N (B) mutants. Electron density maps (2F_0_-F_C_ map, *blue* mesh) and bromine anomalous density maps (*magenta* mesh) are contoured at the 1.5σ and 5σ levels, respectively. Merged structures of E148D *(yellow*, C) and E148N (*gray*, C); wild-type (*yellow*, D) and E148Q (*gray*, D) near Cl^-^ binding sites (*S_ext_* and *S_cen_*). The balls indicate Br^-^ (*magenta)* inEl48N or Cl^-^ (*green*) inEl48Q mutant structure.

**Table 1.** Crystallographic statistics

| Protein, ion | E148D,<br>20 mM Br<sup>-</sup> | E148D,<br>50 mM Br<sup>-</sup> | E148D,<br>200 mM Br<sup>-</sup> | E148N,<br>20 mM Br<sup>-</sup> | E148A,<br>50 mM BrAc<sup>-</sup> |
| --- | --- | --- | --- | --- | --- |
| PDB I.D. | #6AD7 | #6AD8 | #6ADA | #6ADB | #6ADC |
| **Data collection** |  |  |  |  |  |
| Space group | C121 | C121 | C121 | C121 | C121 |
| a, b, c (Å) | 233.2, 97.6,<br>170.8 | 231.4, 100.1,<br>172.6 | 232.4, 100.0,<br>170.8 | 231.9, 98.0,<br>170.8 | 233.2, 96.7,<br>169.8 |
| $\alpha, \beta, \gamma$ (°) | 90, 131.8, 90 | 90, 132.6, 90 | 90, 131.9, 90 | 90, 132.0, 90 | 90, 131.5, 90 |
| Resolution (Å)<sup>a</sup> | 49.0 – 2.95<br>(3.06 – 2.95) | 50.5 – 3.3<br>(3.42 – 3.3) | 50 – 3.15<br>(3.27 – 3.15) | 49.6 – 2.7<br>(2.8 – 2.7) | 49.9 – 3.06<br>(3.16 – 3.06) |
| $R_{merge}$ | 0.06 (0.55) | 0.08 (0.57) | 0.06 (0.57) | 0.12 (1.13) | 0.15 (1.33) |
| $I / \sigma I$ | 6.2 (1.3) | 5.6 (1.2) | 18.1 (2.5) | 34.1 (1.5) | 32.1 (1.6) |
| Completeness (%) | 96.2 (96.7) | 96.6 (98.2) | 99.5 (98.2) | 99.7 (98.1) | 99.4 (99.3) |
| Redundancy | 6.2 (6.0) | 6.3 (6.4) | 6.3 (6.3) | 7.4 (6.7) | 7.5 (7.4) |
| **Refinement** |  |  |  |  |  |
| Resolution (Å) | 49.0 – 2.95 | 50.0 – 3.3 | 34.0 – 3.15 | 34.0 – 2.7 | 34.9 – 3.06 |
| No. reflections | 113927 | 83063 | 98122 | 153235 | 102941 |
| $R_{work} / R_{free}$ | 22.3 / 26.9 | 22.7 / 28.4 | 23.1 / 28.9 | 23.7 / 27.8 | 22.6 / 27.5 |
| No. atoms |  |  |  |  |  |
| Protein / ligand | 13241 / 2 | 13241 / 4 | 13241 / 4 | 13241 / 6 | 13241 / 14 |
| *B*-factor |  |  |  |  |  |
| Protein / ligand | 72.8 / 124.4 | 94.8 / 146.3 | 92.5 / 156.0 | 88.7 / 139.7 | 106.9 / 118.2 |
| R.m.s. deviations |  |  |  |  |  |
| Bond lengths (Å) | 0.01 | 0.01 | 0.01 | 0.01 | 0.01 |
| Bond angles (°) | 1.20 | 1.28 | 1.26 | 1.18 | 1.30 |
| Ramachandran plot<sup>b</sup> | 92.5 / 6.1 / 1.4 | 91.7 / 7.1 / 1.2 | 91.6 / 7.2 / 1.2 | 93.7 / 5.3 / 1.0 | 91.5 / 7.2 / 1.3 |
| Clash score<sup>c</sup> | 18.1 | 13.2 | 13.2 | 17.2 | 13.4 |
| MolProbity score<sup>c</sup> | 2.23 | 2.13 | 2.13 | 2.15 | 2.14 |

<sup>a</sup> numbers in parentheses represent values for the highest resolution shell.<sup>b</sup> percentage of residues in the favored / allowed / disallowed region.<sup>c</sup> Clash and MolProbity scores were calculated as described<sup>42</sup>.

Intriguingly, anomalous Br^-^ signals at the **S_cen_**, the central binding site at which Cl^-^always localizes in wild-type CLC-ec1, vanished in the E148D mutant while the side-chain of Asp148 residue sat near the external binding site. Instead, a strong anomalous Br^-^ signal remained at the *S_int_*, the internal binding site, which has been considered a low-affinity anion binding site^29^ (Fig. 3A). In the E148N mutant structure, three anomalous Br^-^ signals were observed along the Cl^-^ transport pathway, just as in E148Q mutant^16^ (Fig. 3B). However, the rotameric position of the Asn148 residue is very different from that of the Gln148 in E148Q mutant. The side chain of Asn148 takes a rotamer horizontally away from the position of Asp148 inducing ~3.2Å movement of the carboxyl carbon from the center of Cl^-^ transport pathway, whereas the Gln148 side chain vertically rotates ~56^°^ with ~3.6Å translation of the carboxyl carbon from the *S_ext_* (Fig. 3C, D). These distinct side chain positions of Asp148 and Asn148 could reflect a conformational change in the E148D mutant upon protonation of Asp148, as occurs between the wild-type and E148Q mutant. Comparison of the solvent accessibilities of Asn148 and Gln148 residues suggests that the E148D mutant experiences difficulty in delivering a proton to the outside of the protein, and vice versa (Supplementary Fig. 3). This could retard H^+^ transport, followed Cl^-^ slippage, and result in an altered Cl^-^/H^+^ stoichiometry in E148D mutant (Fig. 2C).

### The E148D mutant mimics the conformation of a new intermediate state of the transport cycle

What is the driving force to expel Cl^-^ from the central binding site of E148D mutant while Asp148 is present near the external binding site (Fig. 3A)? To obtain a possible explanation, we measured atomic distances between residues involved in Cl^-^ coordination at the central site in the E148D mutant and wild-type CLC-ec1 structures in the presence and absence of halide ions (Fig. 4 and Supplementary Table 4). In the E148D mutant structure, the carboxyl group of Asp148 slightly moved down to the central Cl^-^ binding site compared to the position of Glu148 in the *Middle* position in the wild-type structure, and atomic distances between carboxyl (Asp148) and hydroxyl oxygens (Ser107 or Tyr445) decreased by 1.6~1.9Å (Fig. 4A, B; Supplementary Fig. 4A). Thus, it is natural to imagine that anions are not able to sit at the central binding site in the E148D mutant in considering the ionic radii of Cl^-^ and Br^-^, ~1.8Å and ~2Å, respectively ^34^. Interestingly, the position of the carboxyl group of Asp148 is almost identical to that of Glu148 carboxylate in the halide-free wild-type CLC-ec1 structures previously solved by two independent groups^33,35^ (Fig. 4C and Supplementary Fig. 4).

**Figure 4.**
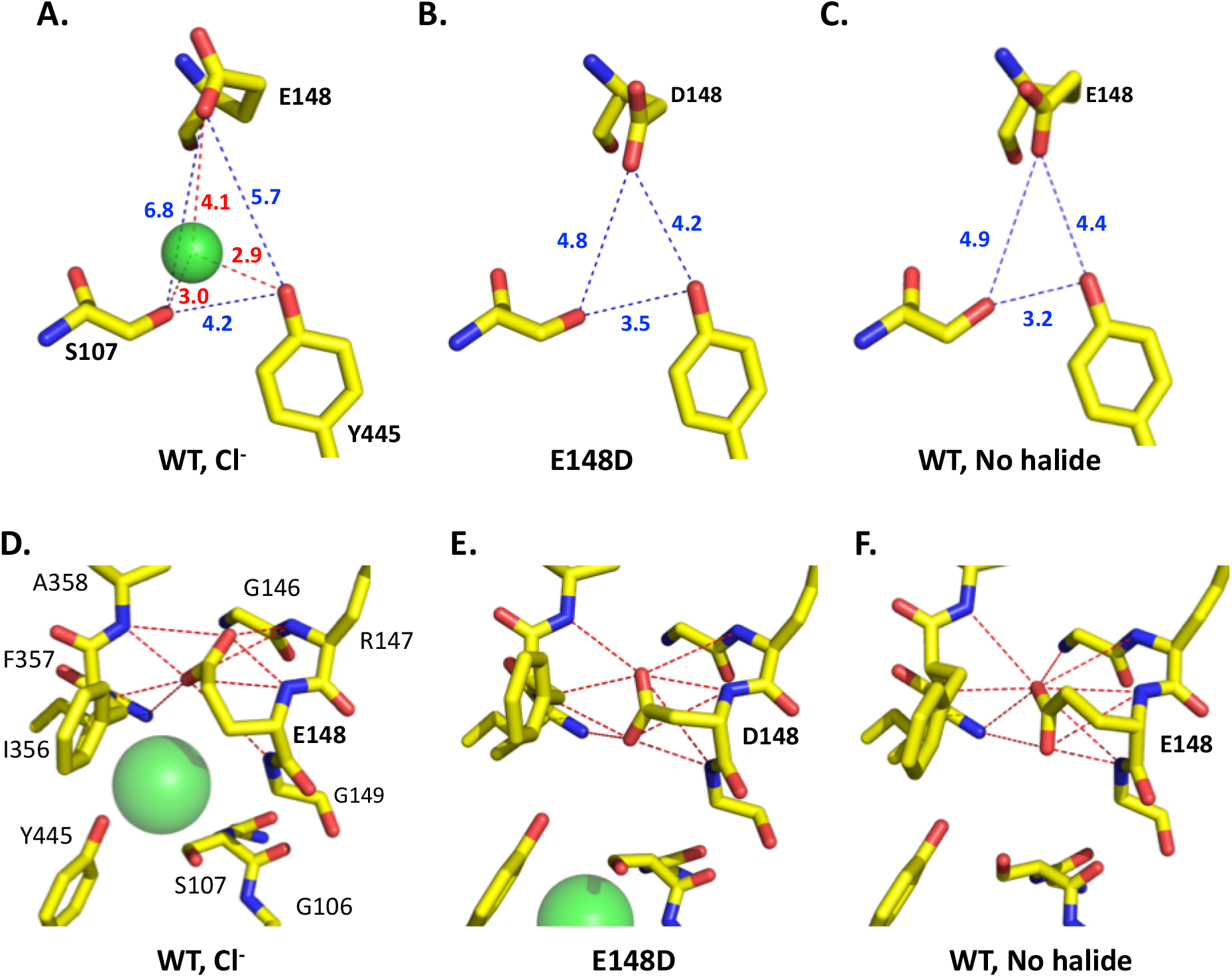
Comparison of atomic coordination between residues involved in anion coordination at the central binding site. Distances among the residues (Glu148, wild-type; Asp148, E148D mutant; Ser107; and Tyr445) involved in Cl^-^ coordination at the central binding site are shown as *blue* dashed lines (A~C). *Red* dashed lines indicate distances between Cl^-^ and coordinating residues. Numbers next to the dashed lines indicate distances in Å. Each distance was measured in chain A from the PDB structures of wild-type and mutant CLC-ec1 (#1OTS for wild-type with Cl^-^ (A), #4KJP for wild-type without halide ion (B), and #6AD7 for E148D (C)) using the PyMOL software. Coordination of neighboring amino acids to the carboxylates of Glu148 and Asp148 residues in wild-type and mutant CLC-ec 1 (D~F). *Red* dashed lines indicate distances less than 4 Å between nearby amino acid residues and γ-carboxylates of the Glu_ex_ of wild-type CLC-ec1 in the presence (D) or absence (F) of Cl^-^, and β-carboxylates of Asp148 in the E148D mutant (E).

In the wild-type CLC-ec1 structure, the backbone amides from helix F and helix N offer an “anion hungry” binding site, *S_ext_*, which Cl^-^ and the deprotonated Glu148 side chain compete with one another to occupy (Fig. 1C) ^15,16^. It is tempting to ask whether the carboxylates of both Asp148 in the E148D mutant and Glu148 in the wild-type in the absence of halide ions have enough coordinating groups to sit in between the external and central binding sites, as Glu148 carboxylate is stabilized by nine backbone amides within a 4Å distance in the wild-type structure while a chloride ion occupies the central binding site (Fig. 4D). The answer is yes: nine to ten backbone amides generate a snug coordination geometry for both cases (Fig. 4E, F). Thus, it can be inferred from the analogy between the structures of halide-free wild-type CLC-ec1 and the E148D mutant that the halide-free conformation is enough to expel Cl^-^ from the **S_cen_**, and that it could be a new intermediate, “*Mid-low*” conformation of Glu_ex_ in the transport cycle.

### The presence of extra anion binding site above the external binding site

A recent study showed that high external Cl^-^ concentration (above 300 mM) significantly slowed both Cl^-^ and H^+^ transports in CLC-ec1 by limiting conformational changes of the Glu_ex_ from *Down-* to *UP*conformations^25^. Also, a computational study on CLC-ec1 suggested an extra Cl-binding site above the Sext^36^. Ün the contrary, it was reported that high Br^-^ concentration up to 400 mM increased anomalous Br^-^ densities at both **S_cen_** and *S_int_* instead of revealing any Br^-^ binding at *S_ext_* or above it in the wild-type CLC-ec1 structure^33^. These data propelled us to identify an anion binding site near *S_ext_* in E148D mutant since Asp148 side chain moves slightly down as a *Mid-low* conformation, which might give a chance to reveal the external anion binding at *S_ext_* or above it.

Impressively, we were able to see an extra anomalous Br^-^ signal above *S_ext_* by increasing Br^-^ concentration: anomalous Br^-^ signal does not appear at the extra binding site (the extra external site, *S_xet_*) in 20 mM NaBr condition, but begins to appear in 50 mM NaBr and becomes prominent in 200 mM NaBr (Fig. 5A; Supplementary Table 3). The bromide ion at *S_xet_* is coordinated by the backbone amide and side chain guanidinium group of Arg147 and is located ~6 Å above *S_ext_* along the anion transport pathway (Fig. 5B). Interestingly a crystallographic water molecule was reported at the position of *Sxet* in the wild-type CLC-ec1 structures^16,28^. These results suggest that *S_xet_* is a genuine low-affinity anion binding site in CLC-ec1, which can be occupied by either a water molecule or halide ion depending on the anion concentration. Thus, it is possible to imagine that the conformational change of Glu_ex_ from *Middle* to *Up*-conformations might be hampered by extra Cl^-^ binding at *S_xet_*, which induces the slowed ion transport in high [Cl^-^] condition as observed previously^25^.

**Figure 5.**
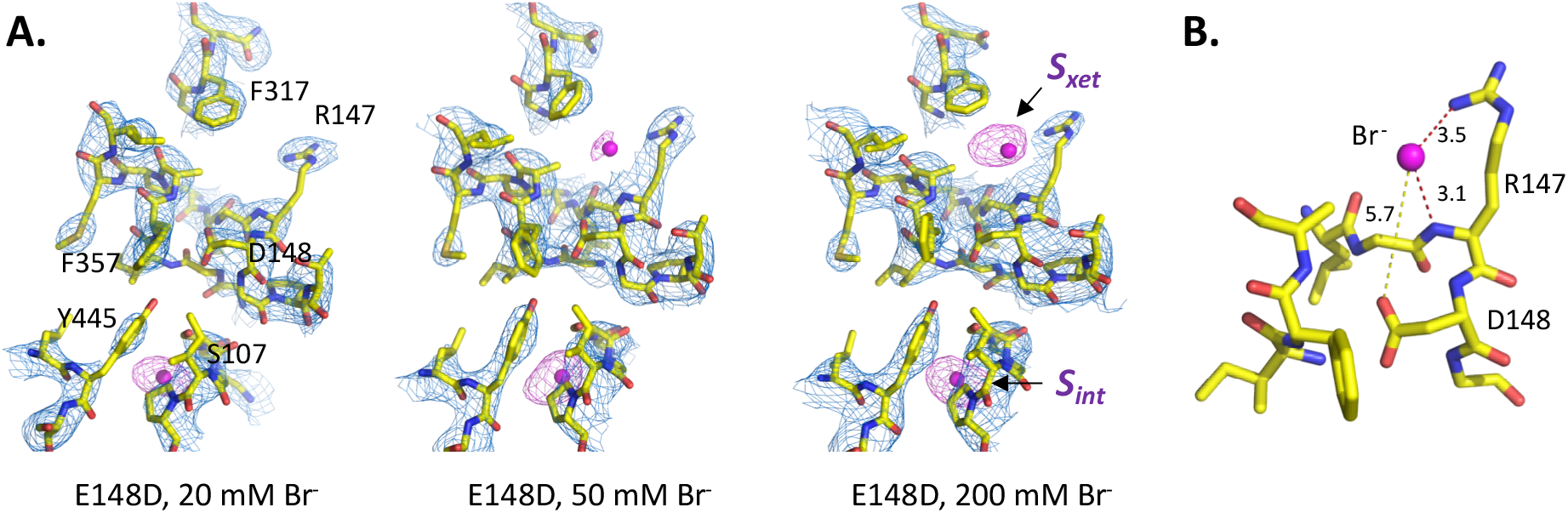
Extra anion binding site above *S_ext_*. A. Structures of E148D CLC-ec1 near anion transport pathway with varied Br^-^ concentration. Electron density maps (2Fo-Fc map, *blue* mesh) and bromine anomalous density maps (*magenta* mesh) are contoured at the 1.5 σ and 4 σ levels, respectively. Anion binding sites are indicated as *S_xet_* (the extra external site) or *S_int_* (the internal site). Atomic distances between Br^-^ at *S_xet_*, and Arg147 or Asp148 residue. Numbers next to the dashed lines indicate distances in Å. Each distance was calculated from chain A in the structures of E148D CLC-ec1 in the presence of 200 mM NaBr.

### Bromoacetate can sit at the S_cen_ in the E148A mutant

Observations of the E148D and wild-type structures indicated that the halide-free CLC-ec1 structure could represent an intermediate conformation of the transport cycle, in which the Glu_ex_ pushes the central Cl^-^ down to the inside as similarly observed in cmCLC^17^. However, whether the conformation of the Glu_ex_ of CLC-ec1 can go further down as it can in cmCLC remains unanswered.

Previously, it was shown that carboxylic acids such as glutamate or gluconate in solution can restore H^+^ transport in the ungated mutant E148A in which the Glu_ex_ is substituted with alanine, and also that glutamate might occupy the **S_cen_** site in E148A CLC-ec1^19^. These observations imply that the Glu_ex_ might go further down in CLC-ec1; however, the presence of glutamate, specifically its γ-carboxylate, at the **S_cen_** is not conclusive, since it was deduced from the omit (*Fo-Fc*) map at a 3.05 Å resolution. Thus, we decided to re-visit a carboxylate occupancy at the **S_cen_** with an anomalously detectable short carboxylic acid, bromoacetic acid (BAA) instead of glutamate (Fig. 6A). Before determining crystal structure of E148A and BAA complex, we first tested whether BAA can bind to the Cl^-^ binding site in E148A CLC-ec1 by measuring the thermodynamic Cl^-^ binding affinities in the presence of a varied concentration of BAA. The binding affinity of E148A CLC-ec1 to Cl^-^ was decreased by ~10-fold (20 μM to 200 μM) in the presence of 10 mM BAA and it was completed abolished by 50 mM BAA (Fig. 6B~D).

**Figure 6.**
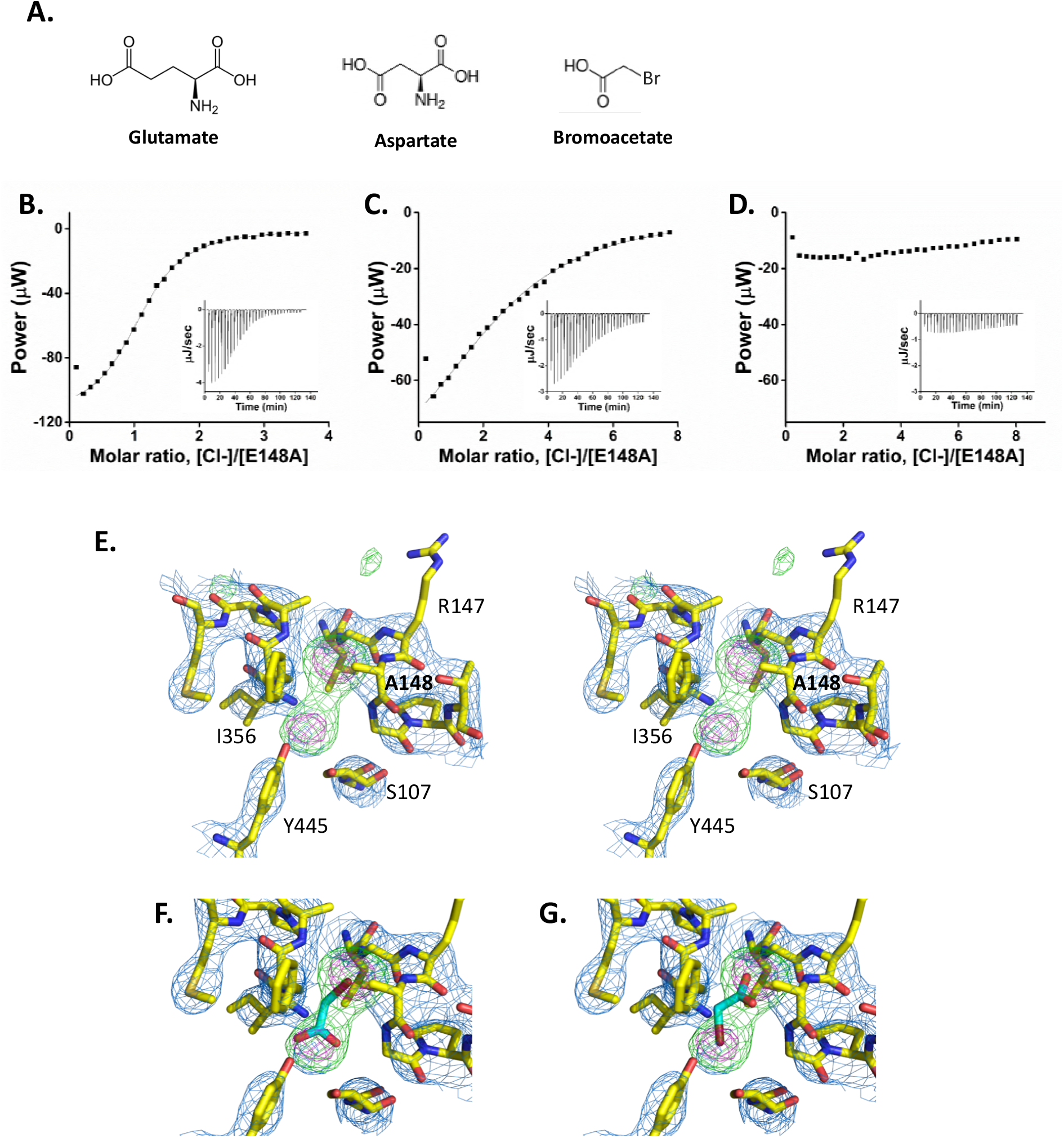
Crystal structure of E148A mutant CLC-ec1 with bromoacetate. A. Chemical structures of glutamate, aspartate, and bromoacetate. Cl^-^ binding affinities of E148A CLC-ec1 in the absence (B) or presence (C, 10 mM; D, 50 mM) of bromoacetic acid in solution. E. Stereo-view structure of the E148A mutant in the presence of 50 mM bromoacetate. Electron density maps (2F_o_-Fc map, *blue* mesh; Fo-Fc positive map, *green* mesh; anomalous bromine density map, *magenta* mesh) are contoured at the 1.5 σ, 4.0 σ, and 12.0 σ levels, respectively. F~G. Alternative conformations of bromoacetate (green) in the Cl^-^ transport pathway.

Interestingly, two anomalous bromine signals were found along the Cl^-^ transport pathway in the E148A mutant structure: a slightly stronger signal near the *S_ext_* and a weaker signal near the **S_cen_** (Fig. 6E; Supplementary Table 3). We speculate that the anomalous signals reflect alternatively positioned bromoacetates in the anion transport pathway, since two anomalous densities are placed in its proximity and *Fo* - *Fc* density is not significantly extruded by the two anomalous signals (Fig. 6E). Based on the anomalous intensities, it can be inferred that both the *carboxylate-down* and *carboxylate-up* configurations are favorable in the Cl^-^ transport pathway. Especially, the bromine group is found at the *S_ext_* and the carboxylate group is placed at the *S_cen_* in the *carboxylate-down* configuration (Fig. 6E~G). These results strongly suggest that the carboxylate of the Glu_ex_ can move down to the *S_cen_* confirming the presence of a cmCLC-like conformation in CLC-ec1. However, this may not necessarily be the case when the side chain is tethered to the protein, since the favorability might depend more on the internal energies of Glu148 rotamers connected to the backbone attached to the protein. Nevertheless, we examined at the chemical properties of carboxylate groups dissociated from the protein in the Cl^-^ transport pathway.

## DISCUSSION

In the present study, we examined a series of Glu_ex_ mutants to gain understanding of the ion transport mechanism of CLC Cl^-^/H^+^ antiporter. From the combined approaches of ion transport measurements, equilibrium binding studies and x-ray crystallography on CLC-ec1, we found the following: (1) aspartate mutation of the Glu_ex_ significantly slows ion transport rate and slightly reduces anion binding; (2) slowed ion transport in the E148D mutant could be attributable to the limited solution accessibility to the Asp148 residue, which makes protonation and deprotonation of the Asp148 side chain difficult, as *S_cen_* in the structure of the E148N mutant; (3) the Asp148 side chain is placed between the *S_ext_* and the *S_cen_*, thus expelling the central Cl^-^; (4) in the absence of halide ions, the Glu148 side chain takes a position identical to that of Asp148 in the E148D mutant; (5) anion can occupy the extra external binding site (*S_xet_*) with low-affinity, which might hinder the conformational change of Glu_ex_ from *Middle* to *Up*; and (6) a short carboxylic acid is able to bind the ungated mutant, E148A at both the *S_ext_* and the *S_cen_* sites.

Üur data provide evidence of intermediate states in the transport cycle of CLC Cl^-^/H^+^ antiporters, and specifically elucidate how movement of Glu_ex_ can be stabilized in the transition between the “*Middle*” and the “*Down*” conformations. The uniquely suggested “*Mid-low*” conformation of the Glu_ex_ is stabilized by backbone amides from helix F and helix N, which are thought to be coordinate the “*Middle*” conformation of Glu_ex_. Once the side chain of Glu_ex_ adopts the “*Mid-low*” conformation, Glu_ex_ can move further to the *S_cen_*, since bromoacetate favors the *carboxylate-down* configuration to the ungated E148A mutant.

The structure of CLC-1 Cl^-^ channel from *Homo sapiens* was recently determined by cryo-electron microscopy^21^. Interestingly, the Glu_gate_ locates to the side of the Cl^-^ transport pathway in CLC-1 while its protonation state is unclear. By coincidence, a notably different rotameric position of Glu_gate_ in CLC-1 is quite similar to that of Asn148 in the E148N mutant (Fig. 3; Supplementary Fig. 5). We examined amino acids propensity at Glu_ex_ position from unique CLC protein sequences to determine whether an Asp variant occurs naturally and were able to find only one case in the infectious bacterium, *Klebsiella pneumoniae* (NCBI accession #WP 102010141.1). Though the Asp substitution at the Glu_ex_ position rarely presents in CLC family proteins, a protonation-dependent rotameric change in the E148D would increase understanding of CLC protein’s transport mechanisms considering that the side chain rotamer of the Glu_ex_ should avoid steric clashes with neighboring amino acids^21^.

Recently, structural studies on a CLC-type F^-^/H^+^ antiporter from *Enterococcus casseliflavus* revealed two conformations, in which the equivalent Glu_ex_ was exposed to either the extracellular face in the *Up* position or to the intracellular face in the *Down* position^22^. Thus, it can be considered a general rotameric twist in CLC-type antiporters that the external glutamate or the gating glutamate switches its positions to *Up, Middle*, and *Down* during the transport cycle regardless of differences in transport mechanism of anion/H^+^ exchanges.

Unlike other alternatively accessing transporters, CLC Cl^-^/H^+^ antiporters have been considered rather static without large conformational changes^37,38^. As a coupled transporter, the Glu_ex_ *up*-conformation, which potentially allows access from both intracellular and extracellular sides, should be avoided to maintain stoichiometrically coupled movements of Cl^-^/H^+^. It has been suggested that both tyrosine (Tyr_cen_) and serine (Ser_cen_) residues near the *S_cen_* act as an inner gate, whose opening occurs only when Glu_ex_ caps the external Cl^-^ entrance^39^. Ün the other hand, a kinetic barrier model explains that a large energy barrier limits rapid movement of Cl^-^ between *S_cen_* and *S_int_* while Glu_ex_ stands in the *up-* conformation^17^. Recent studies have revealed that the CLC-ec1 Cl^-^/H^+^ antiporter undergoes conformational changes of the protein’s backbone: inner gate (Tyr_cen_) opening linked to the movement of helix O^40^ and transition between outward-facing and occluded conformations via rearrangement of helices N and P^41^. However, how the relatively large conformational changes that open either extracellular and intracellular entrances are linked to the rotameric movements of the Glu_ex_ residue warrants further investigation.

## MATERIALS AND METHODS

### Protein purification and crystallography

Analytical-grade chemicals and reagents were purchased from Sigma-Aldrich or otherwise specified, and lipids were from Avanti Polar Lipids. Expression and purification of CLC-ec1 was performed as described^7,28^. The protein was run on a size-exclusion column equilibrated with 100 mM NaCl, 10 mM Tris-HCl, pH 7.5, 5mM Decyl Maltoside (Anatrace) for ion flux experiments.

All crystallographic experiments used ‘ΔNC’ constructs^28^, which is known to be improving the crystal quality of CLC-ec1. For crystallizing CLC-ec1 with Br^-^, proteins were first purified on a cobalt column in low-Cl^-^ buffer (100 mM Na/K tartrate, 2 mM NaCl, 20 mM Tris-SO_4_, 400 mM Imidazole-SO_4_, pH 7.5, 5 mM DM). After removal of the His tag, CLC-ec1-Fab complexes were formed and further purified on a size-exclusion column equilibrated with 20 mM NaBr, 20 mM Na/K tartrate, 10 mM Tris-SO_4_, pH 7.5, 5 mM DM for E148D and E148N mutants, or 50 mM Bromoacetic acid, 10 mM Tris-SO_4_, pH 7.5, 5 mM DM for E148A mutant. Protein was concentrated to 10~20 mg/mL and mixed with an equal volume of crystallization solution in a hanging-drop vapor-diffusion chamber. Crystals, appearing in 20-28% (w/v) PEG 400, 0-20 mM Na/K tartrate, pH 7.5-9.0 in 3-10 days at 22 °C, were cryo-protected by slowly increasing PEG concentration in the mother liquor to ~35%, followed by freezing in liquid N2. Data sets were collected at beamline BL-5C or BL-11C Pohang Accelerator Laboratory (PAL; Pohang, Korea) at X-ray wavelength of 1 Å, or at 0.919 Å for Br^-^ anomalous diffraction. Data were integrated and scaled using HKL2000 or imosflm, and initial models were obtained by molecular replacement against 4ENE or 1OTS using Phaser in the CCP4 software suite. Models were rigid-body refined in REFMAC5 and further refined in Phenix. Protein Data Bank accession codes for each crystal are reported in Table 1.

### Isothermal titration calorimetry

All ITC experiments were performed with a microcalorimeter (Nano ITC, TA Instruments, 300-μL sample volume). For retaining the C−-free conditions, protein was purified on a cobalt column in low-Cl^−^ buffer (100 mM Na/K tartrate, 2 mM NaCl, 20 mM Tris-SO_4_, 400 mM Imidazole-SO_4_, pH 7.5, 5 mM DM), followed by size-exclusion chromatography in no-Cl^-^ buffer (100 mM Na/K tartrate, 10 mM Tris-SO_4_, pH 7.5, 5 mM DM). For Bromoacetic acid (BAA) inhibition experiments, 10~50 mM BAA was added to no-Cl^-^ buffer. Protein (150–250 μM) was titrated with 1.0 – 1.5 μL injections of 2.5 – 40 mM Cl^−^ or Br^−^ at 25 °C. Data were analyzed with single-site isotherms using NanoAnalyze 3.7.5 software.

### Cl^-^ and H^+^ flux assay

Formation of liposomes reconstituted with wild-type or mutant CLC-ec1 (1–5 μg protein/mg lipid) and ion flux measurements have been described in detail^28^. *E. coli* phospholipids were used for all liposome experiments. Briefly, large multilamellar liposomes formed from several freeze-thaw cycles were extruded with a 0.4 mm filter. For H^+^ uptake, a 0.1 mL liposome sample loaded with 300 mM KCl, 40 mM citrate-NaOH, pH 4.8 was passed through a 1.5 mL Sephadex G-50 (GE healthcare) column swollen in 1 mM KCl, 299 mM K-isethionate, 2 mM glutamate-NaOH, pH 4.8, and diluted into 1.8 mL of the same solution in a stirred cell, with pH monitored continuously with a glass electrode. K-isethionate solutions were prepared by titrating isethionic acid (Wako Pure Chemical Industries) with KOH. Cl^-^-driven H^+^ uptake was initiated by addition of 2 mg/mL valinomycin (Vln) and terminated by 2 mg/mL H^+^ ionophore FCCP. Each experiment was calibrated by addition of 50-150 nanomoles of HCl. Cl^-^ efflux was performed similarly except that slightly different buffer systems were used. Liposomes loaded with 300 mM KCl, 25 mM citrate-NaOH, pH 4.5 were diluted as above into 1 mM KCl, 299 mM K^+^-isethionate, 25 mM citrate-NaOH, pH 4.5. For pH-dependent Cl^-^ efflux, Liposomes loaded with 300 mM KCl, 15mM Citrate, 15 mM MES, 15 mM Hepes-NaOH, pH 4.0 ~ 6.0 were also diluted as above into 1mM KCl, 299 mM K-isetionate, 15mM Citrate, 15 mM Mes, 15mM Hepes-NaOH, pH 4.0 ~ 6.0. Efflux of Cl^-^ was triggered by Vln/FCCP, and 30 mM β-octylglucoside was added at the end of the run to determine total trapped Cl^-^. Cl^-^/H^+^ stoichiometry was measured by comparison of initial slopes of H^+^ uptake and Cl^-^ efflux performed in 1–10 mM KCl, 290–300 mM K-isethionate,2 mM Citrate-NaOH, pH 5.2 using liposomes same as in H^+^ uptake experiments.

## Acknowledgments

We thank members of the Lim laboratory for timely help throughout the work, and the staff at beamlines 5C and 11C at PALII (Pohang Light Source II, Pohang Accelerator Laboratory, Pohang, Republic of Korea) for assistance at the synchrotron. We are grateful to Drs. C. Miller (Brandeis University), C.-S. Park (GIST), and J. L. Robertson (University of Iowa) for constructive discussion and criticism on the manuscript. This work was partly supported by KBRI Basic Research Program funded by Ministry of Science and ICT (18-BR-01-02) to H.-H. L, NRF Brain Research Program funded by Ministry of Science and ICT (2017M3C7A 1048086) to H.-H. L, and Korea Health Technology R&D Project funded by the Ministry for Health and Welfare (H18C1254) to H.-H. L.

## Supplementary Figure 1. pH-dependency of E148D mutant

Representative Cl-transport traces for indicated transporters at pH 4.0 (black line) and pH 6.0 (red line). Scales indicate 0.2 unit of relative Cl-concentration in 10 seconds.

## Supplementary Figure 2. Overall structure of E148D and E148N mutants

Merged structure of wildtype (black, pdb #4ENE), E148D mutant (red), and E148N mutant (blue).

## Supplementary Figure 3. Solvent accessibilities of E148D and E148N mutants

Slice view of E148Q (A) or E148N (B) mutant structures. The side chain of Gln148 (A) or Asn148 (B) is represented as spheres. Proton exchange between solution and protein inside is indicated as line and arrow.

## Supplementary Figure 4. Structural comparison of wildtype CLC-ec1 in the presence and absence of halide ions, and E148D mutant

Wildtype (yellow, pdb #1OTS) with Cl-(green sphere),E148D mutant (green), wildtype without halide 1 (cyan, pdb #4KJP), and wildtype without halide 2 (gray, pdb #2EXW).

## Supplementary Figure 5. Structural comparison of human CLC-1 and E148N mutant CLC-ec1 near the Cl-binding sites

E148N mutant (yellow) with Br^-^(*magenta sphere*) and human CLC-1 Clchannel (*cyan*) with Cl^-^ (green sphere, pdb #6COY). Residues are indicated in single-letter symbols with amino acid positions (black, E148N mutant; blue, human CLC-1).

## Supplementary Table 1. Cl-transport metrics

* The values indicate the mean ± the standard errors from 3~6 observations.

## Supplementary Table 2. Thermodynamic values of anion binding at pH 7.5

(a) Adopted from reference 29.

(b) The values indicate the mean ± the standard errors (n>2) or ranges (n=2).

## Supplementary Table 3. Anomalous Br-signal density

* The values indicate the average anomalous peak densities ± the standard errors (n>2) or ranges (n=2).

## Supplementary Table 4. Atomic distances between anion coordinating residues

* The values indicate the average distance ± the standard errors (n>2) or ranges (n-2).

